# Characterization of nanoparticles and fluorescent recombinant extracellular vesicles with three high-sensitivity flow cytometers

**DOI:** 10.64898/2026.02.18.704754

**Authors:** Estefanía Lozano-Andrés, Ye Tian, Sten F.W.M. Libregts, An Hendrix, Xiaomei Yan, Ger J.A. Arkesteijn, Marca H.M. Wauben

**Affiliations:** Division of Cell Biology, Metabolism and Cancer, Department of Biomolecular Health Sciences, Faculty of Veterinary Medicine, Utrecht University, Utrecht, The Netherlands; Division of Infection & Immunology, Department of Biomolecular Health Sciences, Faculty of Veterinary Medicine, Utrecht University, Utrecht, The Netherlands; MOE Key Laboratory of Spectrochemical Analysis & Instrumentation, Key Laboratory for Chemical Biology of Fujian Province, State Key Laboratory of Physical Chemistry of Solid Surfaces, Department of Chemical Biology, College of Chemistry and Chemical Engineering, Xiamen University, Xiamen, Fujian 361005, People’s Republic of China; Laboratory of Experimental Cancer Research, Department of Human Structure and Repair, Ghent University, Ghent, Belgium; Cancer Research Institute Ghent, Ghent, Belgium

**Keywords:** extracellular vesicles, exosomes, high-sensitivity, flow cytometry, silica nanoparticles, characterization, cross-platform, reference material

## Abstract

High-sensitivity flow cytometry (FC) allows multiparametric analysis of nanoparticles (NPs) and extracellular vesicles (EVs). With new instruments available, studies that evaluate their performance using the same materials in a controlled environment are required. Here, we performed a comparative study to investigate the capabilities of three flow cytometers, namely the NanoFCM (NF), BD Influx (IF) and CytoFLEX LX (CF). Firstly, we analyzed a mixed population of silica NPs (SiNPs, 68, 91, 114 and 155 nm) by using light-scatter based detection thresholds (SSC, FSC, VSSC) across a concentration range from 10^6^ to 10^9^ particles/mL. Next, we analyzed fluorescent recombinant EVs (rEVs) by comparing light-scatter based thresholding (488 nm SSC available for all platforms), the combination of SSC thresholding with a fluorescent gate, and fluorescent thresholding for their qualitative and quantitative analysis. We here provide the strengths and limitations for each platform regarding the analysis of differently sized NPs at different sample concentrations.

## Background

Flow cytometry (FC) has become a powerful nanotechnology for high-throughput analysis of single nanoparticles (NPs) and extracellular vesicles (EVs). It allows the qualitative and quantitative characterization of heterogenous cell-derived EV samples as long as the specific requirements are met [1, 2]. Traditionally, the forward scatter (FSC) parameter in FC is used to define the minimum signal required to detect an event, also known as threshold (t). For cells, this threshold is typically set well above the background noise of the instrument. However, in the case of EVs, which are orders of magnitude smaller than cells and scatter considerably less light, the threshold often approaches the instrument’s noise level, posing significant challenges for their reliable detection [3]. In the past decade, diverse FC approaches have been proposed for the characterization of such submicron sized particles [2]. First reports of single EV FC analysis utilized optimized and tailored first generation FC instruments and employed a fluorescence-based detection strategy, requiring all events of interest be sufficiently bright to be detected [4, 5]. A second generation of instruments that collect light scattering with higher sensitivity (violet 405 nm laser side scatter (VSSC)) and accommodate both cellular and small particle measurements, made the use of FC in the EV field more accessible [1, 6]. These second-generation flow cytometers are often equipped with enhanced sensitivity features, e.g., a small particle detector combined with VSSC or the 488nm laser SSC, allowing a single light scatter parameter to be used for EV detection. In parallel, a third generation of instruments with a bottom-up design fully dedicated for nanoparticle analysis was initiated with a state-of-the-art, lab-built nanoflow cytometer. This instrument utilizes an SSC parameter from the 532 nm laser, achieving a sizing resolution comparable to cryo-EM for EV characterization [7] [8]. The technology has since been commercialized by NanoFCM Inc., demonstrating broad utility across diverse applications [9-11].

Despite these advancements, analyzing the whole size range of EVs at the single-particle level remains challenging due to their great heterogeneity, highly variable concentrations and signals that frequently are close to, at and/or below the limit of detection (LOD) of the instrument. Previous studies often used polystyrene (PS) NPs to report the LOD. However, the refractive index (RI) of PS is higher than that expected from EVs, resulting in PSNPs scattering one to two orders of magnitude more light than EVs of a similar size [3]. Alternative materials with a lower RI, such as silica nanoparticles (SiNPs) are thus essential for instrument characterization when unknown samples of low RI will be analyzed. Such SiNPs have been used on dedicated third generation instruments to report size estimations of EVs based on Mie theory modelling [7, 12].

Besides synthetic NPs, reference materials (RM) that are carefully designed to resemble biological EV samples will further aid the validation of flow cytometers able to detect and quantify single particles with high sensitivity [12]. Stable fluorescent biological recombinant EVs (rEVs) and synthetic EV-mimetics composed of a lipid bilayer and canonical EV markers with an average diameter of 125 nm and RI of 1.39 are described [13-15]. Currently, the rEVs developed by Geeurickx et al. are commercially available under the name of exosome standards, fluorescent (Sigma Aldrich) [15].

In this comparative study, we evaluated the detection of mixed populations of SiNPs or PSNPs, and fluorescent rEVs using identical samples and instrument settings across three generations of high-sensitivity flow cytometers: a fully tailored first generation optimized BD Influx™ (IF), a commercial second generation CytoFLEX LX™ (CF) and a state-of-the-art third generation NanoFCM™ (NF). The same samples were measured simultaneously on each platform in the same location by dedicated operators. We here show the influence of particle size and sample concentration on the performance of the different platforms, and identified the limitations of various detection strategies based on light scatter and fluorescence thresholding. Furthermore, we demonstrate the value of well-characterized RM and the use of fluorescent signals for comparing instrument performance and achieving a reliable particle quantification.

## Material & Methods

Silica Nanoparticles (SiNPs). The Silica Nanospheres Cocktail (S16M-Exo, NanoFCM), a mixture of 68, 91, 113, and 155 nm silica nanoparticles (SiNPs), was provided by NanoFCM Inc. The SiNP samples were diluted in Milli-Q H_2_O (filtered 0.22 μm, Merck) prior to the measurements.

Recombinant fluorescent EVs (rEVs). Commercially available fluorescent recombinant EVs expressing green fluorescent protein (GFP) (SAE0193-1VL, Sigma Aldrich). Production and characterization have been previously described in detail [15]. Dilutions to obtain the desired concentrations were made in commercial PBS (Corning) shortly before measurements. The stock concentration used for these experiments was determined by fluorescent NTA to be 1.35E+13 fluorescent particles/mL.

Set up of flow cytometer platforms.

The NanoFCM N30 model (NanoFCM Inc., Xiamen, China), herewith referred to as NF, was equipped with a 488 nm laser line and Single Photon Counting Modules (SPCM) detectors. The fluorescence collection was performed using a 525/40 nm band-pass filter (central wavelength/bandwidth). Low sample flow rate for the measurements was calibrated as 0.005 μl/min.

The jet in air-based BD Influx (BD Biosciences, San Jose, CA), herewith referred to as IF, was equipped with 488, 405, 561 and 635 nm laser lines and PMT detectors were used. An improved reduced-wide angle FSC (rw-FSC) detection was used [16]. Further details about its configuration and adaptations are described elsewhere [4]. The band-pass fluorescence collection filter used was 530/40 nm. Low sample flow rate was calibrated as 6 μl/min.

The CytoFLEX LX (Beckman Coulter, Brea, CA), herewith referred to as CF, with a cuvette-based system was equipped with 375, 405, 488, 561, 638 and 808 nm laser lines and APD detectors. The band-pass fluorescence collection filter used was 525/40 nm. Low sample flow rate was estimated as 10 μl/min.

Acquisition time for every sample was fixed at two minutes for all three platforms. Detailed descriptions of each instrument and methods are provided in Data S1 (MIFlowCyt checklist) and Data S2 (MiFlowCyt-EV framework). Data analysis was performed in FlowJo Version 10.5.0. Data was handled in Microsoft Excel and figures were prepared using GraphPad Prism version 10.0 (GraphPad Software Inc).

## Results

### Light scatter-based thresholding analysis of a mixed population of non-fluorescent silica nanoparticles shows differences in sensitivity across platforms

We first used the SSC detector from the blue laser (488 nm) present on all three instruments to set the threshold as a comparison point due to the absence of a FSC detector on the NF and the lack of sensitivity from the FSC detector on the CF for small particle detection. Firstly, we measured a buffer control (Mili-Q control, here used as dilution buffer) with the selected settings for EV analysis. This buffer control revealed the number and distribution of background signals after a fixed time-based measurement of 120 seconds across all instruments (Figure 1a-c, left graphs). Next, we measured the mixed population of non-fluorescent SiNPs with an increasing diameter size including populations of 68, 91, 114 and 155 nm.

**Figure 1.**
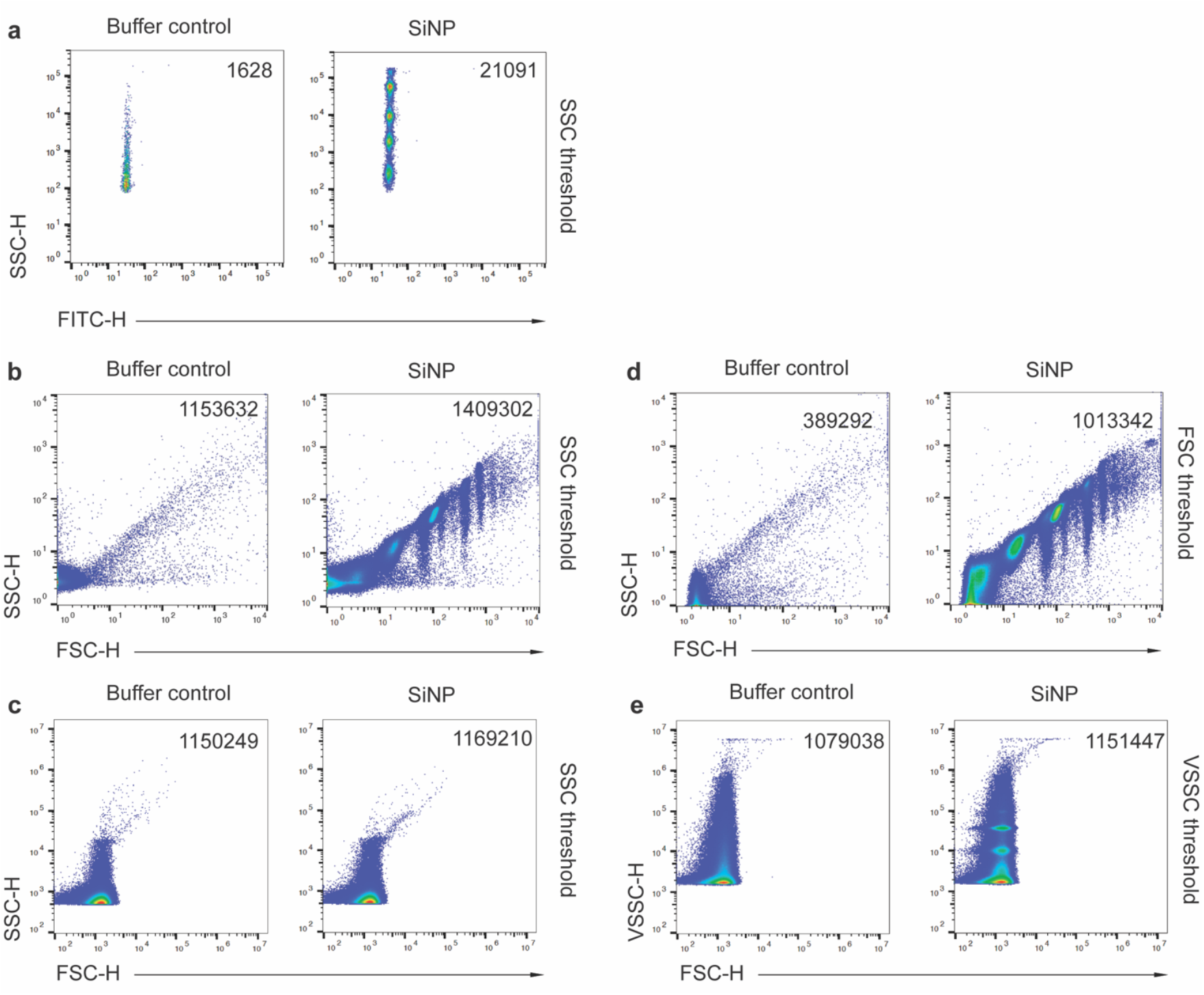
Light scatter-based analysis of buffer control and a mixed population of non-fluorescent silica nanoparticles across platforms. Dot plots displaying number of events as indicated in buffer control and the sample containing 4 sized populations of non-fluorescent silica nanoparticles (68, 91, 114, 155 nm in diameter). Samples were measured on the NanoFCM with **(a)** a SSC (488 nm) threshold, on the BD Influx with **(b)** a SSC (488 nm) and **(d)** a rw-FSC (488 nm) threshold and on the CytoFLEX LX with **(c)** a SSC (488 nm) and **(e)** a VSSC (405 nm) threshold. SiNP samples were pre-diluted shortly before measurements (1:100 for NF and 1:1000 for both IF and CF). Identical buffer control and SiNP samples were acquired simultaneously on all three instruments in the same room under the same conditions.

The NF detected all four populations based on their SSC (488 nm) signals (Figure 1a, right), although the smallest populations were not resolved from the background events detected in the buffer control (Figure 1a, left). The SSC (488 nm) threshold on the IF (IF-SSCt) rendered a lower sensitivity compared to the NF, as the 91 nm SiNP population was the smallest detected population when compared to the buffer control (Figure 1b). Indicating that the 68 nm SiNP population was not discriminated from the background on the IF. When we used the SSC (488 nm) threshold on the CF (CF-SSCt), none of the four SiNP populations were resolved based on SSC and/or FSC signals from the blue laser, and the measured SiNP sample showed events similar to the buffer control (Figure 1c). Alternatively, on both the IF and the CF, we had the possibility to set the threshold based on other light scatter parameters that were more favorable for these instruments. On the IF, we used the reduced wide angle-FSC threshold (488 nm) (IF-FSCt) and on the CF we used the Violet SSC threshold from the 405 nm laser (CF-VSSCt). When using the IF-FSCt we observed 2.9-fold less events in the buffer control (Figure 1d, left) as compared to the SSC (488 nm) threshold (Figure 1b, left), mainly due to a lower background contribution of optoelectronic components. This improved the detection of the 91 nm SiNP population (Figure 1d, right) but the smallest 68 nm SiNP population was still not detectable. The CF-VSSCt measurements showed similar number of events in the buffer control compared to the SSC (488 nm) threshold (Figure 1c and 1e, left panels). However, the VSSC parameter revealed three populations in the SiNP sample (Figure 1e, right panel), with the 91 nm SiNP being the smallest detected population, albeit not separated from the background signals.

To validate that the 91 nm SiNPs were above the LOD on the IF and CF, the 91 nm SiNPs were measured in the absence of the other three sized populations. Compared to the multi-peak sample, the same 91 nm SiNP population was observed consistently across platforms (Figure S1). Lastly, we measured a PSNP sample containing four populations of different sizes ranging from 64, 94, 127 to 156 nm in diameter with the most favorable light scattering thresholding options from the previous dataset (namely NF-SSCt, IF-FSCt and CF-VSSCt). These results revealed that the four populations could be detected on the three platforms, including the smaller 64 nm PSNPs (Figure S2), which is expected due to the higher RI of the polystyrene material compared to silica and subsequently to biological EVs.

### Low and high sample concentration impairs robust analysis of nanoparticle subpopulations across platforms

To investigate the dynamic concentration range on the three different platforms, we prepared six dilutions of the SiNP sample containing 4 sized populations and measured with the most favorable light scattering threshold for each platform (NF-SSCt, IF-FSCt and CF-VSSCt). The most concentrated SiNP sample (1:100 dilution) was favorable on the NF, allowing to clearly visualize the four sized populations (Figure 2a, left). However, lower concentrations (1:4000 and 1:8000 dilutions) showed an impaired discrimination of the four bead populations (Figure 2a, right panels). This can be explained by particles contained in the buffer, becoming more abundant relative to the particles of interest in the sample, thereby compromising the discrimination of the beads overlapping the most with the background (i.e., the 68 nm and 91 nm populations). Importantly, using the NF-SSCt it was possible to perform a robust single particle analysis using a stock concentration of 10^11 particles/mL that was diluted to a measuring concentration of 10^9 particles/mL (1:100 dilution) without suffering from coincidence or swarm detection. In contrast, this high measuring concentration induced swarm detection when measured on both the IF-FSCt and the CF-VSSCt (Figure 2b and c, left panels). The known characteristics of swarm detection, i.e., non-linear relationship between the number of detected events and the dilution factor, and aberrant light scattering signals [17, 18], were clearly visible in the 1:100 and 1:500 diluted sample acquired with CF-VSSCt (Figure 2c) and to a lower extend in the 1:100 diluted sample acquired with IF-FSCt (Figure 2b). Hence, the data from these concentrated samples obtained with the CF and IF do not provide reliable information on single particle detection. Collectively, our results show that low particle concentrations challenge single particle detection on the NF, while high particle concentrations compromise single particle detection on the IF and CF when light-scatter based measurements are performed.

**Figure 2.**
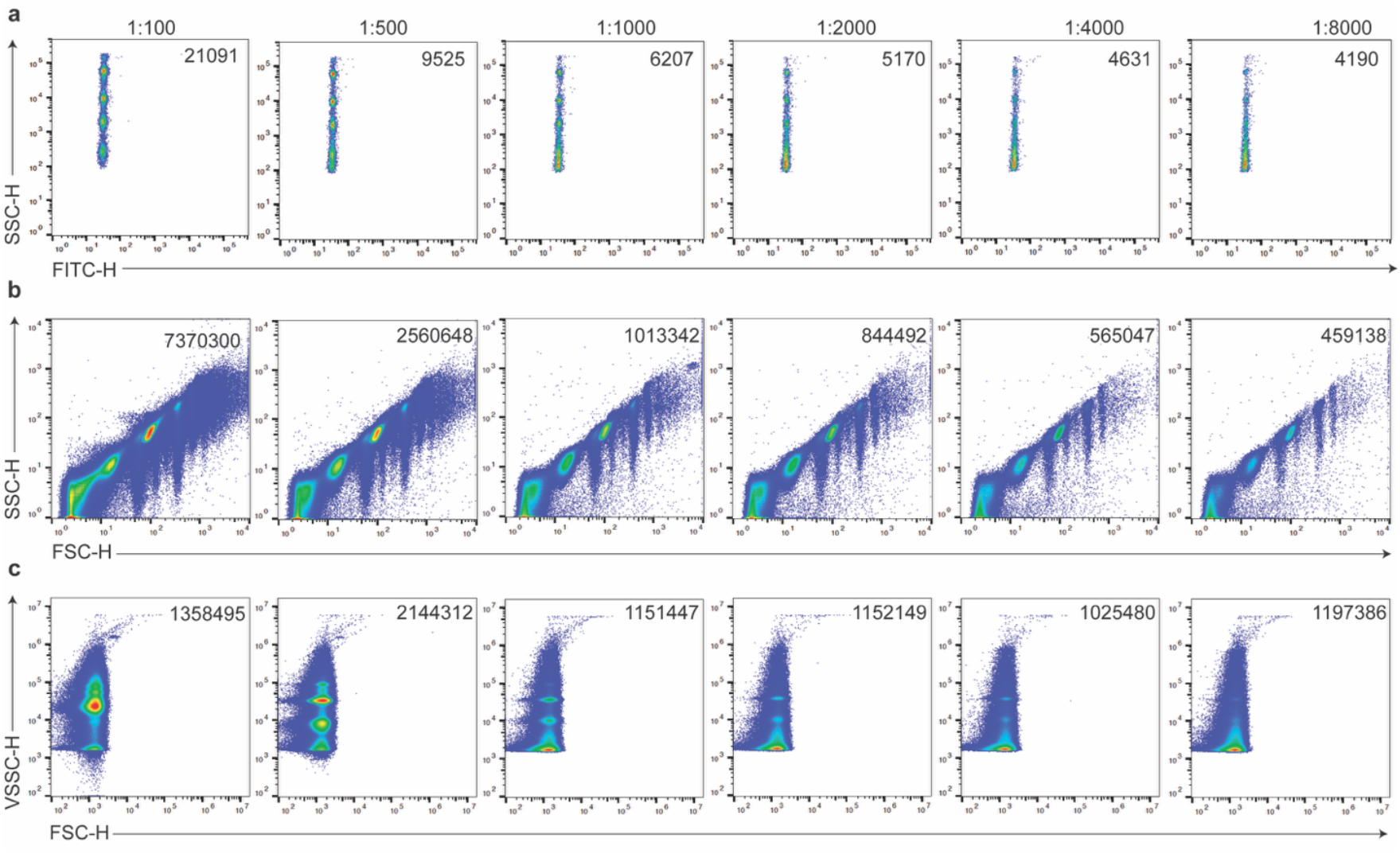
Analysis of six sample dilutions from non-fluorescent silica nanoparticles across platforms. Dot plots displaying SSC-H Vs FITC-H or FSC-H from total number of events of non-fluorescent SiNP measured for 2 minutes **(a)** on the NanoFCM with a SSC (488 nm) threshold **(b)** on the BD Influx with a rw-FSC (488 nm) and **(c)** on the CytoFLEX LX with a VSSC (405 nm) threshold. Samples were diluted as indicated in the top row (from 1:100, 1:500, 1:1000, 1:2000, 1:4000 and 1:8000, left to right).

### Qualitative and quantitative analysis of fluorescent rEVs reveals differences across detection strategies and platforms

To compare the analysis of biological EVs across the three platforms, we conducted parallel measurements using identical samples of commercially available fluorescent rEVs. These rEVs have consistent biophysicochemical properties similar to those of native EVs and contain encapsulated EGFP (enhanced green fluorescent protein). The EGFP allows for its detection in combination with light scattering signals, adding an extra layer of specificity that enables the discrimination of fluorescent rEVs from other particles that might be present in the preparation and the dilution buffers. Firstly, an SSC (488 nm) threshold was applied on all three platforms. Besides the rEV sample, a PBS buffer control, here used as dilution buffer, was measured under the same settings (Figure 3, left panels). Using the NF-SSCt, fluorescent rEVs are resolved as a separate population based on the fluorescence collected in the FITC channel (Figure 3a), with an increased event rate of 12-fold, from 10 ev/s in the buffer control to 120 ev/s in the rEV sample. The IF-SSCt measurements revealed a clear population of consistently increasing light scatter and fluorescence intensities (Figure 3b, top panel), with an increased event rate of 2.6-fold, from 7,483 ev/s in the buffer control to 19,464 ev/s in the sample. Furthermore, for the IF a secondary light scatter parameter, the rw-FSC, could be plotted Vs fluorescence, which revealed a better separation of the fluorescent rEVs from the background (Figure 3b, bottom panel). The CF-SSCt measurements again showed a high number of events in the buffer control (Figure 3d), which translated in a minimal event rate difference between the buffer control (9,827 ev/s) and the rEV sample (10,307 ev/s). Furthermore, the fluorescent rEV were also very difficult to observe in the SSC Vs. Fluorescence dot plot. However,, when the more favorable VSSC parameter for the CF was plotted Vs Fluorescence the fluorescent rEV population was visible (Figure 3d, bottom panel).

**Figure 3.**
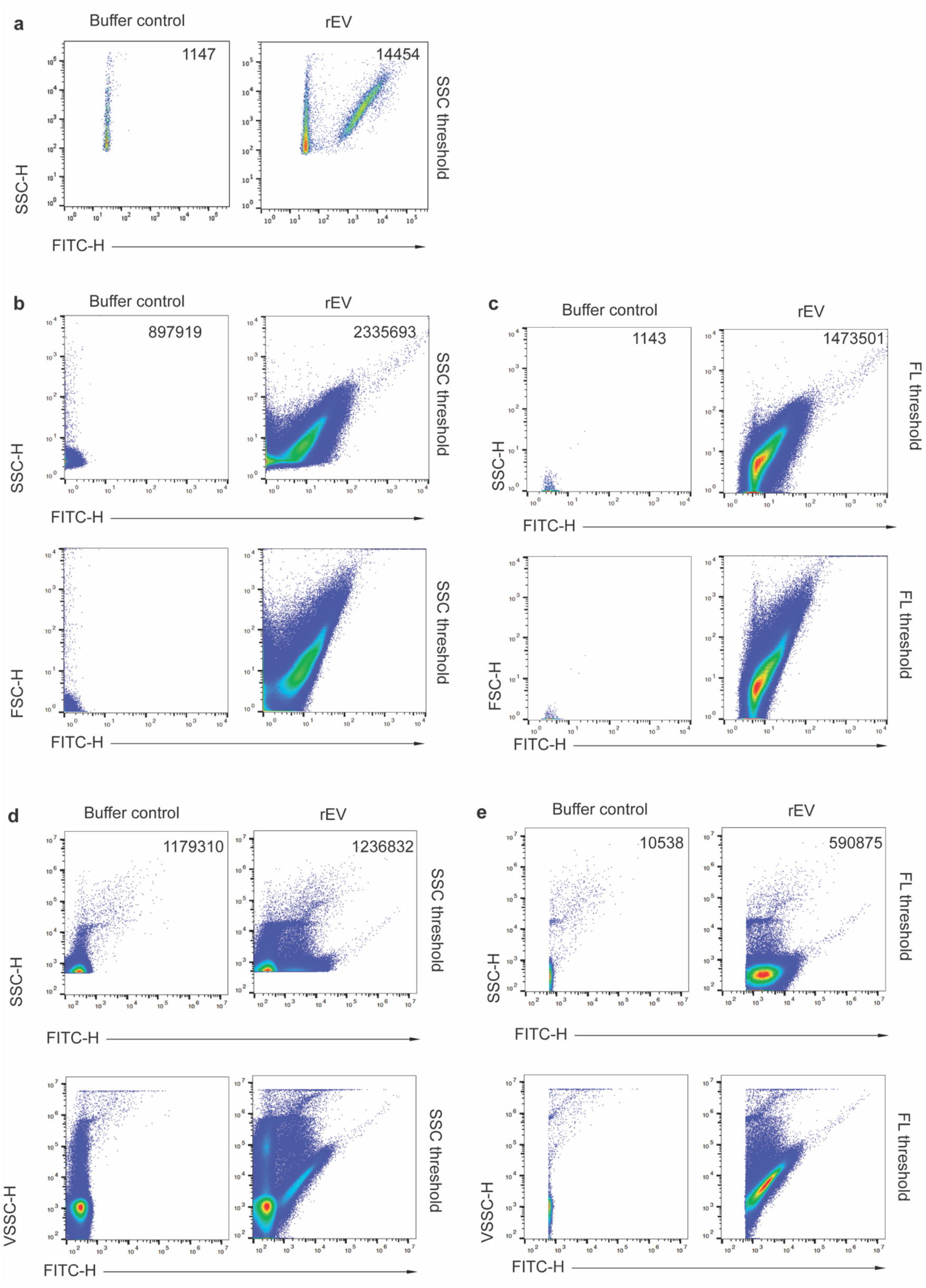
Qualitative analysis of fluorescent rEV across different platforms. Dot plots from the PBS buffer control, here used as dilution buffer, (left) and fluorescent rEVs (right). Measurements **(a)** on the NanoFCM with a SSC threshold, **(b)** on the BD Influx with a SSC threshold and **(c)** a FL threshold, **(d)** on the CytoFLEX LX with a SSC threshold and **(e)** a FL threshold. rEV samples were pre-diluted before measurements (1:1000 for NF and 1:32000 for both IF and CF).

Next, we utilized the EGFP signal present in the rEV to explore fluorescence-based thresholding on the IF and the CF. Since on the NF the fluorescent rEVs are fully separated from the background (Figure 3a), a simple gate suffice to select the fluorescent population on the NF. In contrast, both IF and CF require to set the threshold prior acquisition and recording of the samples. Using this strategy, background events in the buffer control were strongly decreased as expected, resulting in 785-fold and 111-fold reduction respectively for IF-FLt and CF-FLt measurements (Figure 3c and 3e, left panels). This strategy also proved to not compromise the detection of the fluorescent population of rEV (Figure 3c and 3e, right panels), which was followed by the measurement of consecutive sample dilutions (Figure S3).

Since EV concentration is the most reported metric in literature utilizing single particle EV analysis and has been proven relevant in health Vs disease conditions, we next calculated the particle concentrations of measured rEVs with either SSCt, SSCt and fluorescence gating (SSCt+FLg) (gating strategies are shown in Figure S4), or FLt. Due to the previously observed differences in sample concentration that yielded an optimal event rate across the three instruments measurements of the fluorescent rEV were performed across six reciprocal dilutions of the sample (1:1000, 1:2000, 1:4000, 1:8000, 1:16000 and 1:32000). Since the NF fully resolved the fluorescent rEV population using SSCt, fluorescent events were gated from the SSCt measurements. To correct for the observed differences between instruments in terms of events present in the dilution buffer we subtracted those from the sample and normalized to the dilution factor and flow rate. Remarkably, we found that the NF-SSCt detected rEV concentrations ranging from 1E12 to 1E13 events/mL after correcting for the dilution factor, flow rate and buffer control background. The last is an overestimation by the NF of the actual number of rEVs present in the most diluted sample when using a SSCt (Figure 4). Overall, the measurements of the most concentrated rEV samples (dilutions 1:1000, 1:2000 and 1:4000) were found more appropriate in terms of background to sample events, while more diluted samples (1:8000, 1:16000 and 1:32000) lead to an increasing overestimation of the rEV concentration using the NF-SSCt (Figure 4). Quantification using the three most concentrated rEV samples (1:1000, 1:2000 and 1:4000) revealed 1.7E12 events/mL ± 27% (rSD). This variability was effectively reduced when a fluorescent gating was applied around the EGFP signal present in the rEV population (NF-SSCt+FLg), resulting in an average concentration of 7E11 events/mL ± 4% (rSD). These findings illustrate that robust quantitative measurements can be achieved when EV samples are analyzed in an appropriate dilution and that fluorescence signals ensure reliable detection of particles of interest.

**Figure 4.**
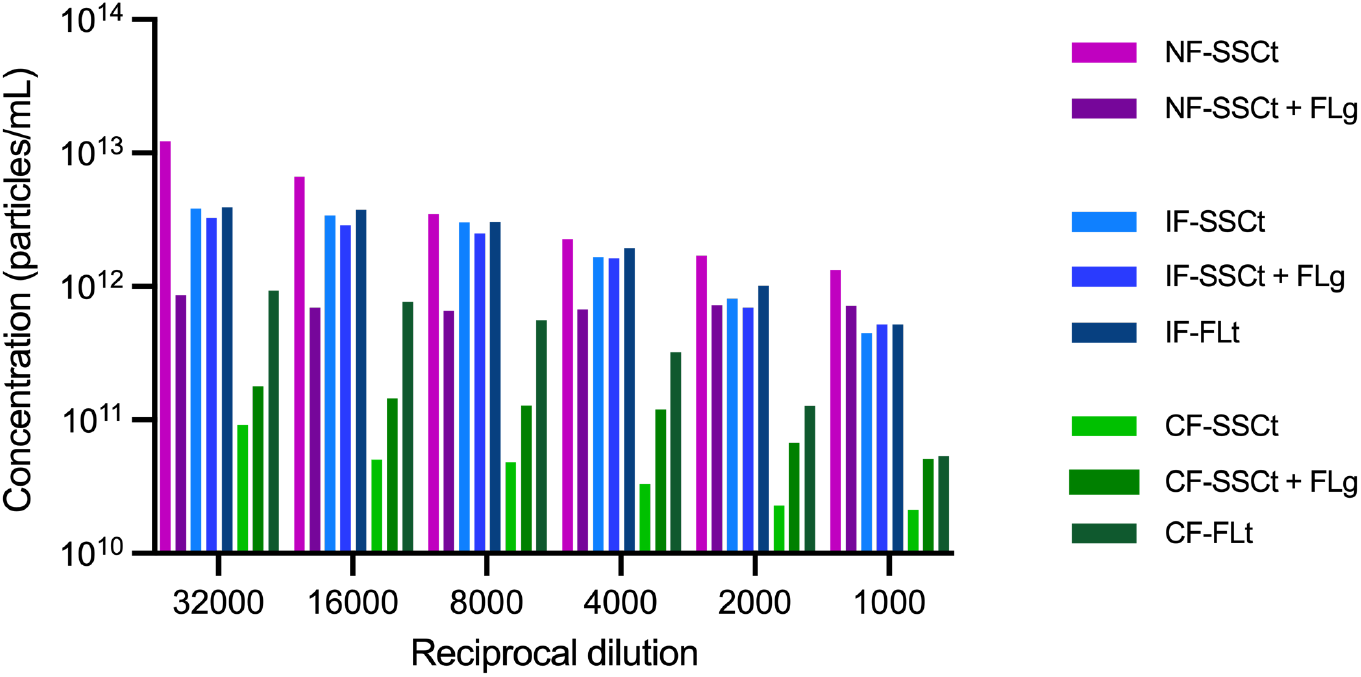
Quantitative analysis of fluorescent rEV. Bar graph of measured rEV concentrations using the indicated dilution across and different threshold options (SSCt or FLt). For comparison, we used a fluorescent gate (FLg) over the total events detected with SSCt to select for the fluorescent rEV population on the NanoFCM (NF), BD Influx (IF) and CytoFLEX LX (CF). Concentration measurements are corrected for time, dilution factor, PBS control and flow rate for each instrument and detection mode.

The IF-SSCt, IF-SSCt+FLg and IF-FLt calculated rEV concentrations ranged from 5E11 to 4E12 events/mL for all detection strategies (Figure 4), with the lowest concentrated rEV samples (1:8000, 1:16000 and 1:32000) being identified as suitable for single particle detection. Utilizing these latter three dilutions, the average concentrations of rEVs were 3.4E12 events /mL ± 12% (rSD), 2.9E12 events /mL ± 13% (rSD) and 3.6E12 events /mL ± 13% (rSD) respectively for IF-SSCt, IF-SSCt+FLg and IF-FLt, showing great consistency across the different detection strategies on the IF.

In contrast, the CF measured rEV concentrations showed a bigger discrepancy among the different detection strategies and dilutions, ranging from 2E10 to 9E11 events/mL with the relatively low rEV concentrations mainly caused by the high background noise on the CF Overall, samples with high rEV concentrations (dilutions 1:1000, 1:2000 and 1:4000) caused strong swarm effects, resulting in a drop in events, leading to erroneous quantification and underestimation of the number of fluorescent rEVs present in the sample. Applying fluorescent gating on the CF around the EGFP-expressing rEVs (SSCt+FLg) improved the detection resulting in a higher particle concentration, albeit to a lower extend when compared to directly applying a FLt (Figure 4). Measurements using the CF-SSCt, CF-SSCt+FLg and CF-FLt with the lowest concentrated rEV samples (1:8000, 1:16000 and 1:32000) resulted in an average concentration of 6.4E10 events/mL ±39% (rSD), 1.5E11 events/mL ±17% (rSD) and 7.5E11 events/mL ± 24% (rSD), respectively. From which the last detection strategy (CF-FLt) showed the highest agreement with the other two instruments (NF-SSCt+FLg and IF-SSCt+FLg and IF-FLt). Overall, rEV quantification based on SSC-thresholding only resulted in high differences among instruments, compared to using FL-based detection (either by NF-SSCt+FLg or IF-FLt and CF-FLt). Furthermore, selecting the appropriate sample dilution per instrument is a crucial step for robust EV quantification and fluorescence-based detection revealed the highest quantification agreement among instruments.

## Discussion

The advancement of high-sensitivity FC has opened new opportunities for the multiparametric characterization of heterogeneous samples containing submicron sized particles, including NPs and EVs. To evaluate the strengths and limitations of different instruments, it is important to perform cross-platform studies that measure identical samples and validate the utility of reference materials. Here, we evaluated three instruments from different generations that have reported capabilities for measuring EVs at the single particle level [4, 6, 9, 16]. Previously, the added value of calibration using commercial materials such as fluorescence reference beads with molecules of equivalent soluble fluorophore (MESF) units and non-fluorescent NIST-traceable beads for both light-scatter and fluorescence flow cytometric EV analysis has been clearly indicated and specific modelling software, i.e. FCM PASS, has been developed [3, 19]. However, using these calibration strategies for the comparison of instruments from different generations is not so straightforward. The older generation of instruments with analogue detection, such as the IF used in this study, cannot be calibrated in this way due to the lack of precise signal digitalization [20]. Furthermore, the light scatter detection range of the NF does not allow to measure PS particles bigger than 500 nm in size with the settings required for small EV analysis, thereby hampering the utilization of the available tools to be implemented for studies such as the one presented here. Despite the limitations to implement such advanced standardization methods across all three platforms, we followed the MIFlowCyt-EV framework for data reporting and included the use of relevant controls and well-characterized reference materials (SiNPs and rEVs) [2]. To minimize external variations due to, e.g. storage and transportation of samples [21], we performed the experiments with identical samples simultaneously in the same location. Furthermore, the three platforms were operated in parallel, each by an expert for the respective platform.

To compare light scatter-based measurements across platforms, we analyzed mixed populations of well-characterized SiNPs and PSNPs. While the NF was the only platform that could detect the smallest population of 68 nm non-fluorescent SiNPs, our results show that by low sample concentrations the detection of the smallest SiNP populations (68 and 91 nm) by the NF was impaired due to the overlap with background noise. Both the IF and CF were able to detect the second smallest SiNP population (91 nm), but they required a 10-fold lower sample concentration compared to the NF to ensure single particle detection, which partially accounts for differences in the sample flow rate among the three instruments. Importantly, the high background signals of the CF obscured SiNP analysis to a much greater extend compared to the IF. The later having much lower background interference and two high-sensitivity light scatter parameters. Overall, the background noise can substantially compromise light-scatter analysis of NPs and EVs, indicating a preference for instruments with low background signal when small particles are measured.

Furthermore, we here demonstrate clear differences in the preferred light-scatter triggering strategy between the three platforms, i.e. NF-SSCt, IF-FSCt and CF-VSSCt showing the best discrimination between background signals present in the dilution buffer and NP samples. This illustrates that there is no ‘‘one-fits-all’’ strategy that can be applied to all flow cytometers, but that the choice depends on the design and configuration of the different platforms, as well as the samples that are being measured. Here, we selected a SSC thresholding strategy (448 nm) for comparison purposes across platforms, as it was the only light scatter-based thresholding option available on all platforms. FSC thresholding (488 nm) was only possible on the IF, as the NF completely lacks the FSC and the CF lacks the required FSC sensitivity. Likewise, VSSC thresholding (405 nm) was only available on the CF. Interestingly, our measurements revealed that the selected SSC thresholding for all three platforms exhibited sufficient light scatter sensitivity to detect SiNPs as small as 91 nm and to discriminate subpopulations that differ by only 23 nm in size (114 nm vs. 91 nm). These results show the power of high-sensitivity flow cytometry- and the potential to enable simultaneous subset analysis of other synthetic and biological particles, helping to elucidate sample heterogeneity. Moreover, here we show the added value of mixed reference SiNP populations to investigate the performance of high-sensitivity flow cytometers in relation to single EV analysis. Since, silica has a much lower RI compared to polystyrene, silica is preferred when low scattering particles, such as EVs, will be analyzed [3]. Such mixed reference SiNP populations (up to 500 nm in size) are also of great interest for calibration and standardization of other orthogonal single NP and EV analysis technologies [22].

Besides synthetic NPs, tailored biological reference materials have been proposed as helpful tools to assess instrument performance in relation to EV analysis. Here, we took advantage of commercially available fluorescent recombinant EVs [15] and showed that these rEVs with an average size of 125 nm could be detected on all three platforms, thereby providing a valuable tool for instrument assessment [15]. In addition, using fluorescent rEVs we were able to evaluate their detection by not only using light scatter thresholding but also fluorescence thresholding, which effectively reduced background noise. Moreover, combining light scatter thresholding with fluorescence analysis of these rEVs enabled us to circumvent the impact of undesired background signals, thereby improving the robustness of rEV quantification.

Importantly, we here demonstrate that sample concentration severely impacts both qualitative and quantitative analysis of fluorescent rEVs across flow cytometers. Consistent with our SiNP data, we found that the NF with a 0.005 μL/min sample flow rate required at least a 10-fold higher sample concentration (E9 particles/mL) compared to the IF, with a 6 uL/min sample flow rate and the CF, with a 10 uL/min sample flow rate (E8 particles/mL). We showed that low concentrated samples can be measured on the NF, but that SSC-based detection resulted in accumulation of background events, causing an overestimation of the rEV concentration. On the other hand, the higher rEV concentrations, optimal for the NF, introduced swarm detection (massive coincidence) on the IF and CF, thereby compromising quantitative analysis [1, 17, 18, 23]. Based on these findings, we want to point out that for biological EV samples of unknown concentration, sample dilutions are indispensable and need to be adjusted and validated for each instrument [2]. Besides the reference synthetic NPs that are commonly used for instrument assessment, we therefore advocate the use of well-characterized fluorescent rEVs as a suitable reference material to evaluate and compare EV detection capabilities across different platforms, allowing to investigate both light scatter and fluorescence-based detection. To further advance cross-platform studies and ultimately ease the translation potential of EVs into the clinic, future developments are needed to expand the current portfolio of calibrators in the nanometric range, aiming to be compatible across instruments from different generations, including the newer third generation of fully dedicated high-sensitivity analyzers [24] [25].

## Supporting information

Sup Lozano-Andres E et al sup info

## Acknowledgments

The authors would like to thank Dr. Dimitri Aubert (NanoFCM Co., Ltd, United Kingdom) for great assistance to set-up the NanoFCM instrument in our lab. We thank the Laura Varela and the Flow Cytometry and Cell Sorting Facility of The Faculty of Veterinary Medicine at Utrecht University for support. E.L.A. is ISAC SRL Emerging Leader (2025-2028).

## Author Contributions

E.L.A. designed and performed experiments, analyzed data and wrote the manuscript. Y.T. performed experiments and analyzed data. S.F.W.M.L. performed experiments. A.H. prepared materials. X.Y. gave technical and conceptual advice. G.J.A.A. and M.H.M.W. supervised the research, designed (performed) experiments and wrote the manuscript. All authors critically reviewed and edited the manuscript.

## Data availability

Flow cytometric data files will be made available upon request.

