## Supplementary material for "Characterization of nanoparticles and fluorescent recombinant extracellular vesicles with three high-sensitivity flow cytometers": Sup Lozano-Andres E et al sup info

### Supplementary Material & Methods

**Polystyrene Nanoparticles.** Polystyrene Nanoparticles (PSNPs) (64, 94, 127 and 156 nm in diameter) were purchased from Bangs Laboratories, Inc. These PSNPs were prepared into a mixture and diluted with Milli-Q $H_2O$ (filtered 0.22 µm, Merck) water prior to the measurements.

**Supplementary Table 1. Author Checklist: MIFlowCyt-Compliant Items.**

| Requirement | Please Include Requested Information |
|---|---|
| 1.1. Purpose | The purpose of this study is to evaluate the performance of three different flow cytometers when measuring silica nanoparticles (SiNP) and recombinant fluorescent Extracellular Vesicles (rEV) |
| 1.2. Keywords | extracellular vesicles, exosomes, high-sensitivity, flow cytometry, silica nanoparticles, characterization, cross-platform, reference material |
| 1.3. Experiment variables | Sample dilution (as indicated) and detection strategies |
| 1.4. Organization name and address | Department of Biomolecular Health Sciences<br>Faculty of Veterinary Sciences<br>Utrecht University<br>Yalelaan 2, 3584 CM<br>Utrecht, The Netherlands |
| 1.5. Primary contact name and email address | Dr. Estefanía Lozano Andrés<br> |
| 1.6. Date or time period of experiment | January 2019 |
| 1.7. Conclusions | We provide information on the performance of three |

| | |
|---|---|
| | different high-sensitivity flow cytometers to characterize nanoparticles and EVs |
| 1.8. Quality control measures | Dilution buffer control<br>Serial dilutions |
| 2.1.1.1. (2.1.2.1., 2.1.3.1.) Sample description | Silica Nanoparticles and rEV |
| 2.1.1.2. Biological sample source description | Described under material & methods |
| 2.1.1.3. Biological sample source organism description | Human |
| 2.1.2.2. Environmental sample location | |
| 2.3. Sample treatment description | N/A |
| 2.4. Fluorescence reagent(s) description | N/A |
| 3.1. Instrument manufacturer | NanoFCM, Becton Dickinson and Beckman Coulter |
| 3.2. Instrument model | NanoFCM N30 model<br>BD Influx™ optimized instrument for detection of sub-micron sized particles as described previously.<br>CytoFLEX LX |
| 3.3. Instrument configuration and settings | BD Influx optimized to measure small particles. All configuration details can be found in van der Vlist, E. J., Nolte-'t Hoen, E. N., Stoorvogel, W., Arkesteijn, G. J., and Wauben, M. H. (2012) Fluorescent labeling of nano-sized vesicles released by cells and subsequent quantitative and qualitative analysis by high-resolution flow cytometry. Nat Protoc 7, 1311-1326 and Arkesteijn, G.J.A., et al., Improved Flow Cytometric Light Scatter Detection of Submicron-Sized Particles by Reduction of Optical Background Signals. Cytometry A, 2020. **97**(6): p. 610-619.<br>Briefly, samples were measured at a constant flow rate for 120 seconds using indicated settings (SSC, rw-FSC or FL). |
| 4.1. List-mode data files | Are available upon request. |
| 4.2. Compensation description | No compensation was required since only one fluorescence signal was read. |
| 4.3. Data transformation details | No data transformation was applied. |
| 4.4.1. Gate description | Gates were applied based on the controls (shown in Supplementary Material & Methods). |
| 4.4.2. Gate statistics | The number of total events recorded in 120 seconds measurements are shown in the dot plots. |
| 4.4.3. Gate boundaries | Images of the ungated and gated plots can be found in figures and supplementary data. |

**Supplementary Table 2. MIFlowCyt-EV framework.**

| | |
|---|---|
| 1.1 Preanalytical variables conforming to MISEV guidelines | Yes, all relevant data can be found in provided references. |
| 1.2 Experimental design according to MIFlowCyt guidelines | Yes, MIFlowCyt checklist can be found as part of the supporting information of this manuscript. |

| | |
|---|---|
| 2.1 Sample staining details | Sample staining was not performed |
| 2.2 Sample washing details | No sample washing was performed |
| 2.3 Sample dilution details | Yes, described in the text and indicated in the figures. |
| 3.1 Buffer-only controls | Yes, relevant buffer controls were measure and are shown. |
| 3.2 Buffer with reagent controls | Not required as there was no sample staining performed |
| 3.3 Unstained controls | Not required as there was no sample staining performed |
| 3.4 Isotype controls | Not required as there was no sample staining performed |
| 3.5 Single-stained controls | Not required as there was no sample staining performed |
| 3.6 Procedural controls | Not required as there was no sample staining performed |
| 3.7 Serial dilutions | Yes, serial dilutions were performed and are shown for SiNP and rEV samples. See Figure 2 and S3. |
| 3.8 Detergent-treated controls | Detergent treatment was not performed |
| 4.1 Trigger channel(s) and threshold(s) | On the NF we used a SSC threshold (488 nm laser) with 40 mW and 10% of SSC decay, following the manufacturers instructions.<br>On the IF we used a SSC threshold (488 nm laser at 0.32, a.u.), rw-FSC (488 nm laser at 0.29, a.u.) and FL (488 nm laser at 0.61, a.u.).<br>On the CytoFLEX LX we used a SSC threshold (488 nm laser at 500, a.u.), VSSC threshold (405 nm laser at 1600 a.u.) and FL threshold (488 nm laser at 600 a.u.) |
| 4.2 Flow rate / volumetric quantification | Yes, low flow rate was kept constant across settings for each instrument. Flow rate on the NF was 0.005 ul/min, on the IF was 6 uL/min and on the CF was 10 ul/min. |
| 4.3 Fluorescence calibration | Not performed due to lack of materials available compatible with all instruments at the time of the study |
| 4.4 Scatter calibration | N/A |
| 5.1 EV diameter/surface area/volume approximation | N/A |
| 5.2 EV refractive index approximation | N/A |
| 5.3 EV epitope number approximation | N/A |
| 6.1 Completion of MIFlowCyt checklist | Yes, see Table S1 |
| 6.2 Calibrated channel detection range | N/A |
| 6.3 EV number/concentration | Yes, see Figure 4. |
| 6.4 EV brightness | N/A |
| 7.1 Sharing of data to a public repository | All data files are available upon request |

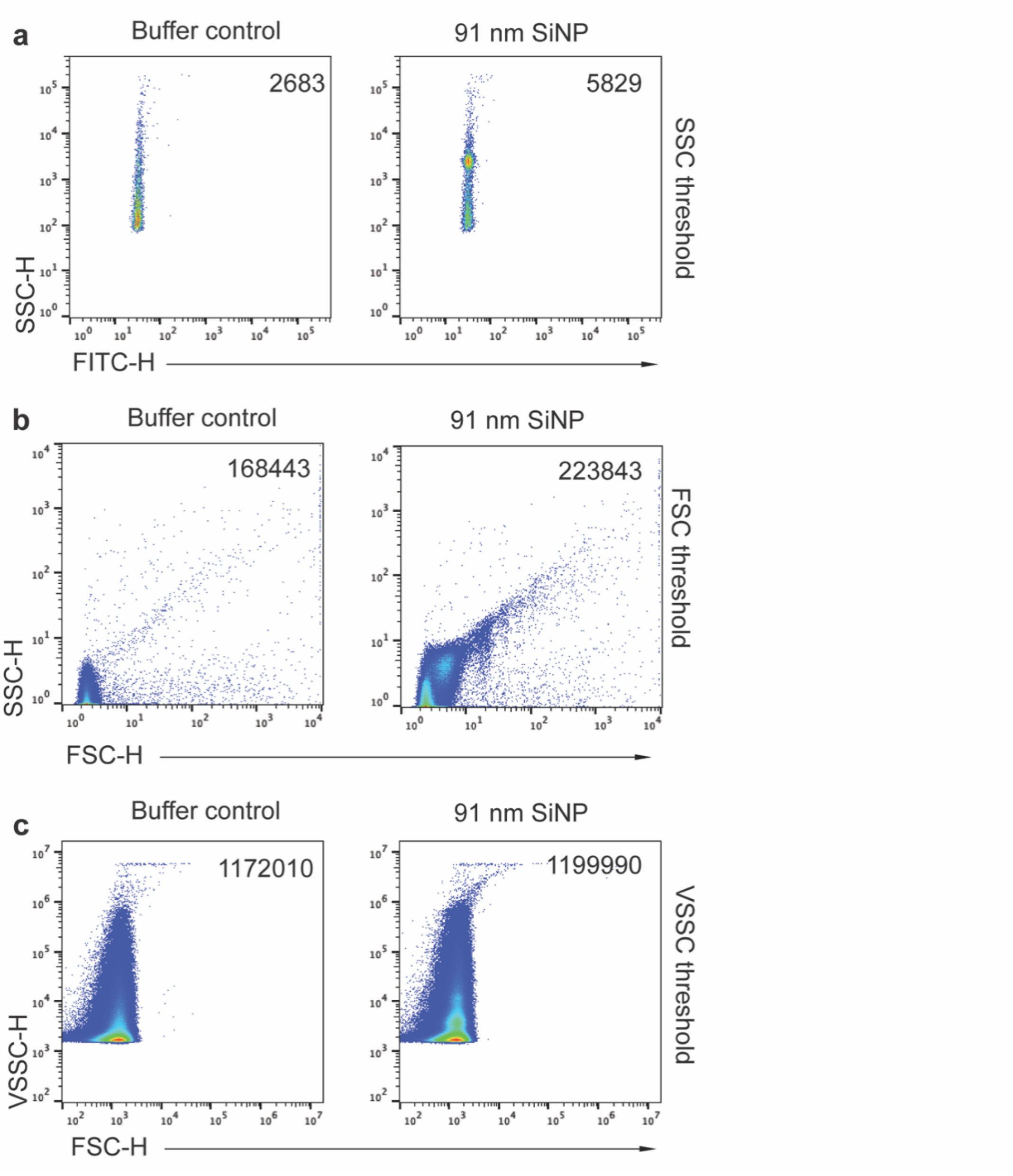


**Figure S1.** Buffer control and Silica nanoparticles of 91 nm diameter were measured across instruments with the selected threshold combinations on (a) NF-SSCt, (b) IF-FSCt and (c) CF-VSSCt.

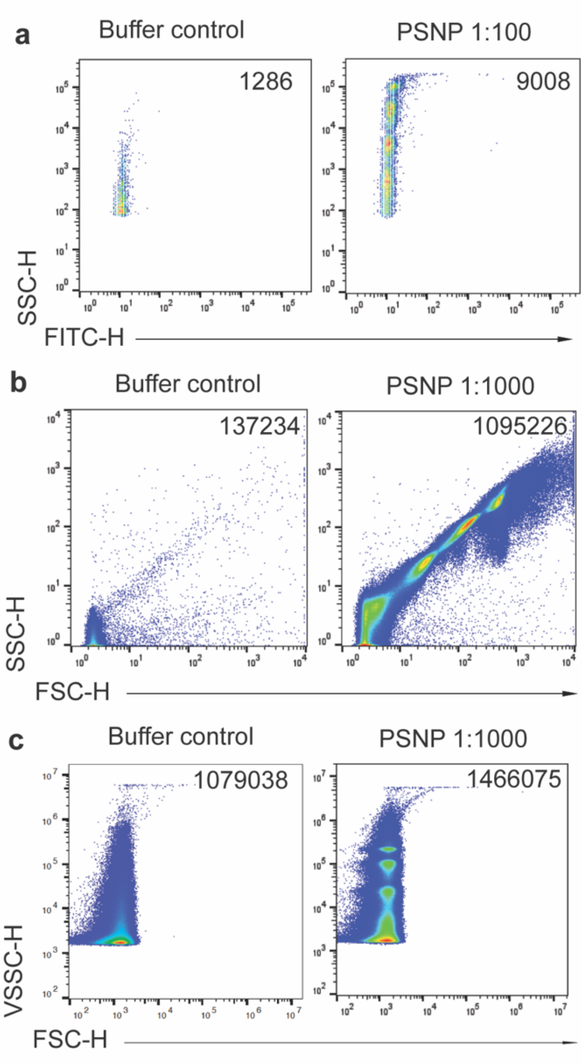


**Figure S2.** Light scatter-based analysis of buffer control and a mixed population of non-fluorescent polystyrene nanoparticles across platforms. Dot plots displaying number of events as indicated in buffer control and the sample containing 4 sized populations of non-fluorescent polystyrene nanoparticles (64, 94, 127 and 156 nm in diameter). Samples were measured on the NanoFCM with (a) a SSC (488 nm) threshold, on the BD Influx with (b) a rw-FSC (488 nm) threshold and on the CytoFLEX LX with (c) a VSSC (405 nm) threshold. PSNP samples were pre-diluted shortly before measurements (1:100 for NF and 1:1000 for both IF and CF). Identical buffer control and PSNP samples were acquired simultaneously on all three instruments in the same room under the same conditions.

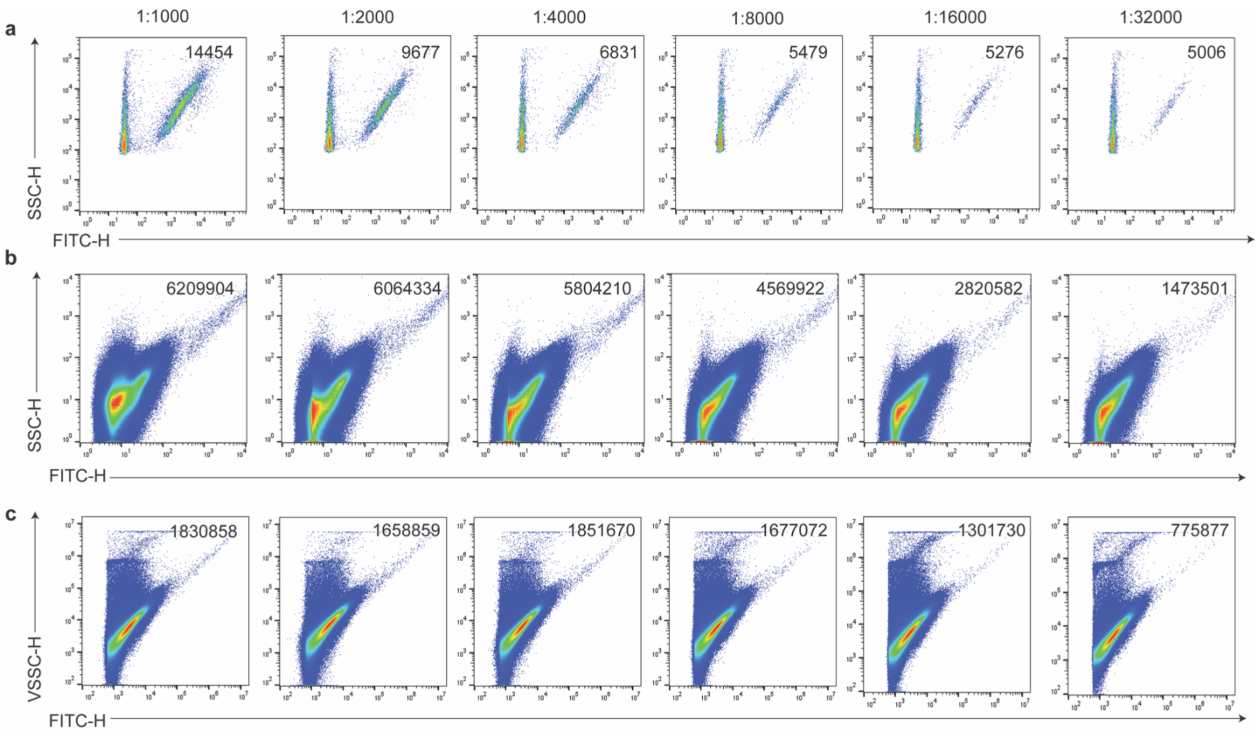


**Figure S3.** Serial dilutions to define single EV detection. Six diluted samples are measured on the (a) NF-SSCt, (b) IF-FLt and (c) CF-FLt. Number of detected events is shown in the top of each dot plot.

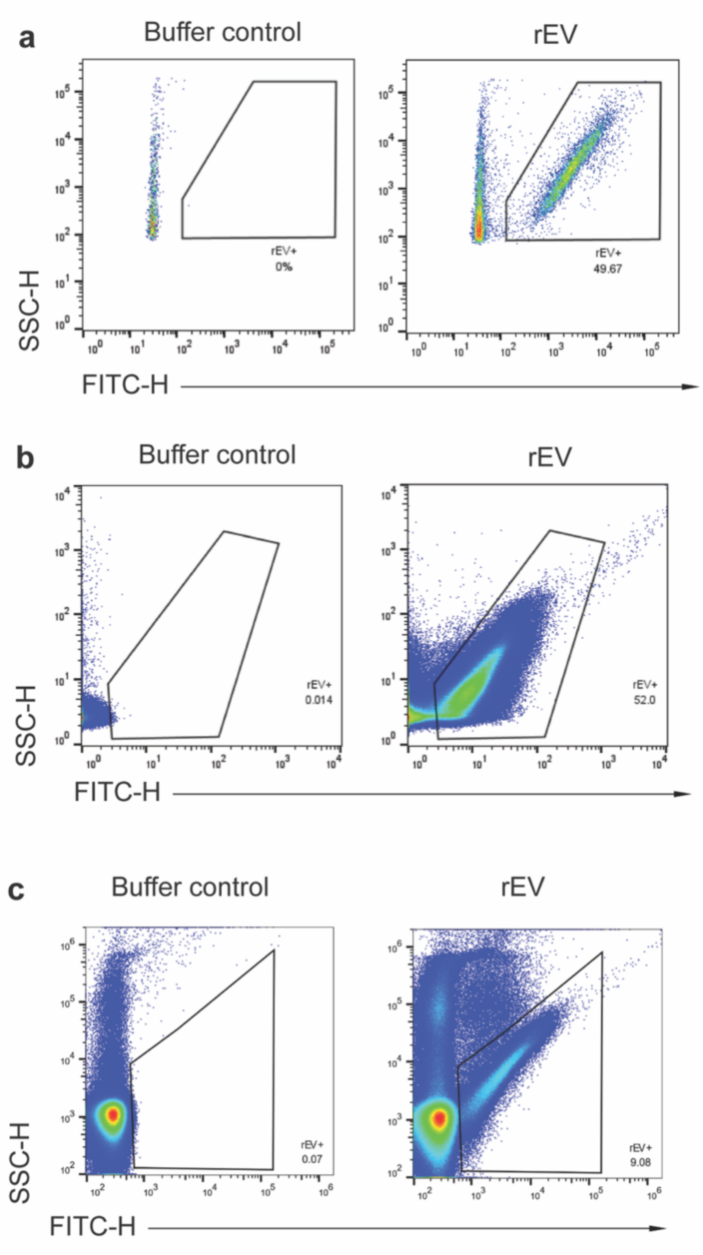


**Figure S4.** Gating strategy for fluorescent rEV samples. Dot plots showing the gated fluorescence (FITC-H) Vs SSC-H on the (a) NF, (b) IF and (c) CF from a buffer control and the rEV sample at dilutions of 1/1000 on the NF, and 1/32000 on both the IF and CF.
